# *Nanjinganthus*: An Unexpected Flower from the Jurassic of China

**DOI:** 10.1101/240226

**Authors:** Qiang Fu, José Bienvenido Diez, Mike Pole, Manuel García-Ávila, Zhong-Jian Liu, Hang Chu, Yemao Hou, Pengfei Yin, Guo-Qiang Zhang, Kaihe Du, Xin Wang

**Author notes:** Corresponding author contact information: **Zhong-Jian Liu**,; Senior author, **Xin Wang**.

## Abstract

The origin of angiosperms has been the focus of intensive botanical debate for well over a century. The great diversity of angiosperms in the Early Cretaceous makes the Jurassic rather expected to elucidate the origin of angiosperm. Former reports of early angiosperms are frequently based on a single specimen, making many conclusions tentative. Here, based on observations of 284 individual flowers preserved on 28 slabs in various states and orientations, we describe a fossil flower, *Nanjinganthus dendrostyla* gen. et sp. nov., from the South Xiangshan Formation (Early Jurassic) of China. The large number of specimens and various preservations allows us to give an evidenced interpretation of the flower. The complete enclosure of ovules in *Nanjinganthus* is fulfilled by a combination of an invaginated and ovarian roof. Characterized by its actinomorphic flower with a dendroid style, cup-form receptacle, and angio-ovuly, *Nanjinganthus* is a *bona fide* angiosperm from the Jurassic. *Nanjinganthus* re-confirms the existence of Jurassic angiosperms and provides first-hand raw data for new analyses on the origin and history of angiosperms.

## Introduction

Despite the importance of angiosperms, a controversy remains as to when and how this group came into existence. Since the time of Darwin, some scholars have proposed that angiosperms existed before the Cretaceous (Smith *et al*. 2010; Clarke *et al*. 2011; Zeng *et al*. 2014; Buggs 2017), although others have favored a Cretaceous origin (Scott *et al*. 1960; Herendeen *et al*. 2017). Such uncertainties make the phylogeny and systematics of angiosperms tentative. Some reports of early angiosperms (i.e., *Monetianthus* (Friis *et al*. 2001)) are based on a single specimen and are thus difficult to test and confirm. Better and more specimens of indisputable age are preferred to test related hypotheses. Here, we report an unusual actinomorphic flower, *Nanjinganthus* gen. nov., from the Lower Jurassic based on the observations of 284 individual flowers preserved on 28 slabs in various orientations and states (Table S1). The abundance of specimens allowed us to dissect some of them and thus demonstrate and recognize a cup-form receptacle, ovarian roof, and enclosed ovules in *Nanjinganthus*. These features, in addition to actinomorphic organization, indicate that *Nanjinganthus* is a true flower of an angiosperm. The origin of angiosperms has long been an academic “headache”, but we think that *Nanjinganthus* will shed a new light on the subject.

## Results

### Genus *Nanjinganthus* gen. nov

**Generic diagnosis**: Flowers subtended by bracts. Bracts fused basally. Flowers pedicellate, probably bisexual, actinomorphic, including four whorls of organs, epigynous, with inferior ovary. Sepals 4–5, more or less rounded in shape, with two lateral rib-free wings, attached to the receptacle rim with their whole bases, each with about four longitudinal ribs in the center, surrounding the petals when immature but reflexing when mature, with epidermal cells with straight cell walls. Petals 4–5, cuneate, concave, and with two lateral rib-free wings, with rounded tips and about four longitudinal ribs in the center, surrounding the gynoecium when immature but reflexing when mature, with epidermal cells with straight cell walls. Possible stamen including a stalk and four pollen sacs. Gynoecium in the center, unilocular, secluded by a cup-form receptacle from the bottom and sides and by an integral ovarian roof from the above. Style centrally located on the top of the ovarian roof, dendroid-formed. At least two ovules inside the ovary, elongated oval, pendulous and inserted on the ovarian wall by a thin funiculus, with the micropyle-like depression almost opposite the chalaza.

**Type species**: *Nanjinganthus dendrostyla* gen. et sp. nov.

**Etymology**: *Nanjing-* for Nanjing, the city where the specimens were discovered, and -*anthus* for “flower” in Latin.

**Type locality**: Wugui Hill, Sheshan Town, Xixia District, Nanjing, China (N32°08′19″, E118°58′20″) (Figs. S1).

**Horizon**: The South Xiangshan Formation, the Lower Jurassic.

### Species *Nanjinganthus dendrostyla* gen. et sp. nov

(Figs. 1–5; Figs. S4h, S5-S10)

**Figure 1.**
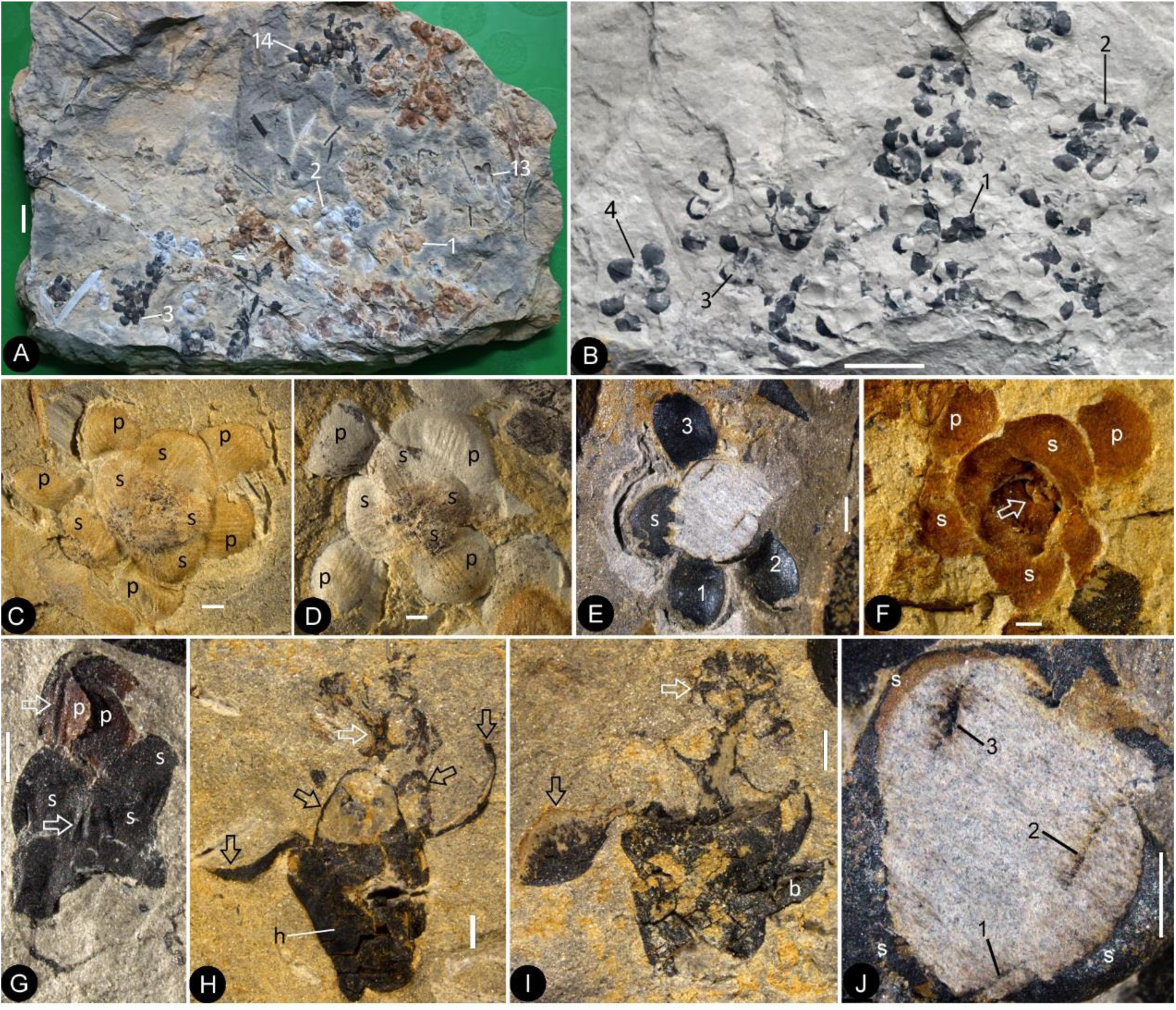
Flowers preserved in different states and their details. Bar = 1 mm except otherwise annotated. **A**. Numerous flowers preserved on a single slab. The numbered ones are detailed in the later figures. PB22222B. Bar =1 cm. **B**. Numerous coalified flowers on the same slab. The numbered ones are detailed in later figures. The possible stamen in Fig. 4i is from flower 4. PB22223. Bar = 1 cm. **C**. Bottom view of Flower 1 in Fig. 1a, showing 5 sepals (s) and 5 petals (p) with longitudinal ribs. PB22222B. **D**. Bottom view of Flower 2 in Fig. 1a, showing 4 sepals (s) and 4 petals (p) with longitudinal ribs. PB22222B. **E**. Bottom view of the flower in Fig. S6f, showing a sepal (s) and three petals (p) radiating from the center, which is broken to show the relationship among the sepals and petals as in Fig. 1j. PB22278. **F**. Top view of Flower 1 in Fig. S5b with sepals (s), petals (p), and seeds (arrow, enlarged in Fig. 3h) inside the receptacle. PB22226. **G**. Side view of a flower bud (No. 1 in Fig. 1b) with longitudinal ribs (arrows) on the sepals (s) and petals (p). PB22223. **H**. Side view of Flower 1 in Fig. S5d, showing a receptacle (h), petals and sepals (black arrows), and a dendroid style (white arrow). PB22224. **I**. Side view of Flower 2 in Fig. S5d, showing petals (black arrow), the dendroid style (white arrow), and the scale (b) on the receptacle. PB22224. **J**. Detailed view of an oblique section of the flower shown in Fig. 1e, showing the arrangement of three petal bases (1–3) inside the sepals (s). These petals bases correspond to the three petals (1–3) in Fig. 1e.

**Figure 2.**
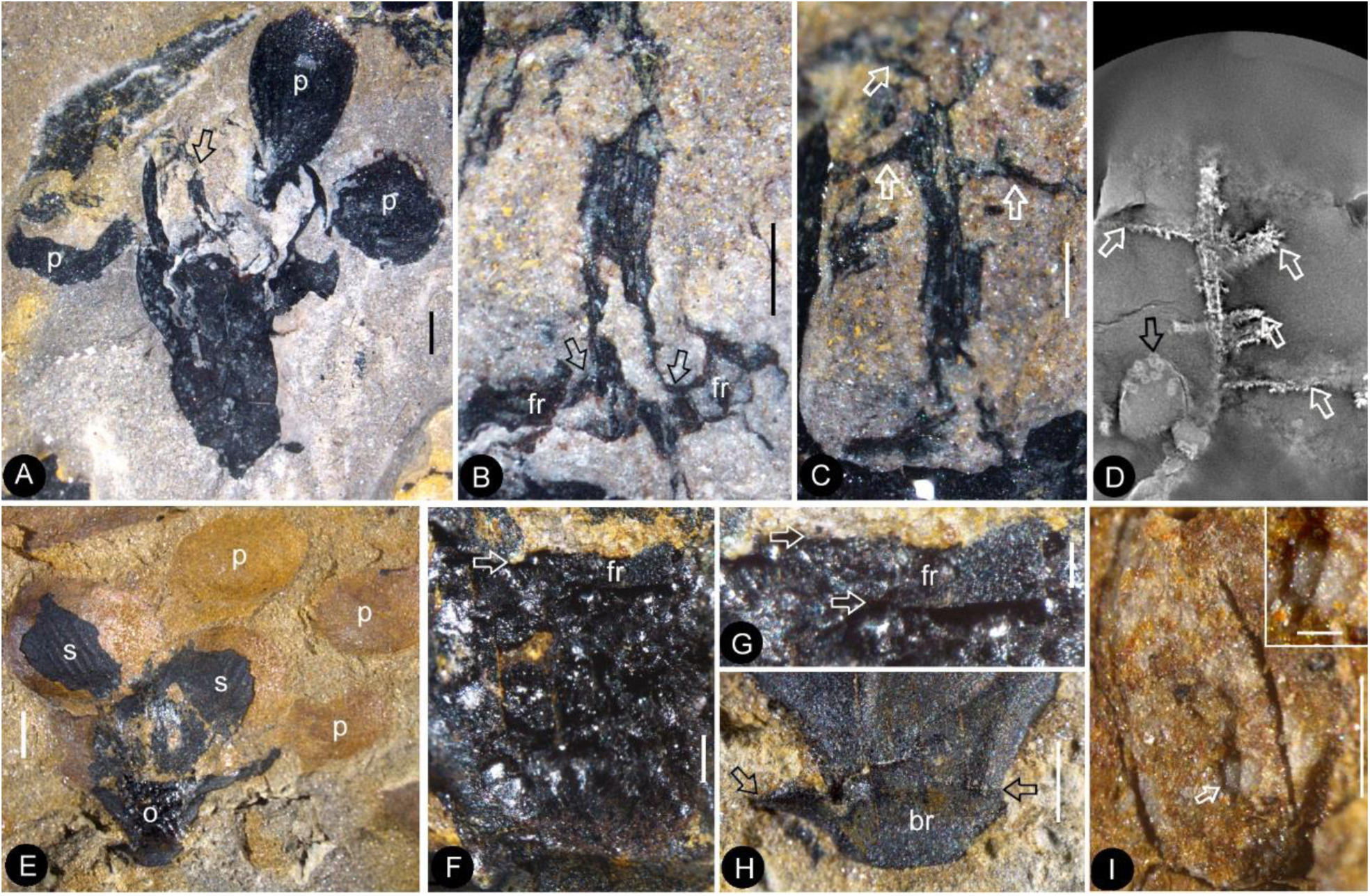
The flowers and their internal details. Bar = 1 mm except otherwise annotated. **A**. A flower carefully dégaged to expose the details of the gynoecium. Note the petals (p) and a style (arrow) in the center. PB22282. **B**. Detailed view of the style in Fig. 2a, showing its connection (arrows) to the ovarian roof (fr). Bar = 0.5 mm. **C**. Distal portion of the same style as in Fig. 2b, showing its dendroid form with lateral branches (arrows). Bar = 0.5 mm. **D**. Micro-CL slice 1169 showing a sepal/petal (black arrow) and branches (white arrows) of the style, embedded in sediments, of flower 4 in Fig. S5e. PB22222a. **E**. Side view of an organically-preserved flower with sepals (s) and petals (p). Note the dark organic material in the ovary (o) and some sepals. The foreground portion of the receptacle has been removed to show the internal details in Figs. 2f-h. PB22281. **F**. Detailed view of the receptacle/ovary in Fig. 2e. Note the ovarian roof (fr) preventing the outside (above) sediment (yellow color) from entering the ovarian locule. Bar = 0.2 mm. **G**. Detailed view of the solid organically-preserved ovarian roof (fr) with integral outer (upper arrow) and inner (lower arrow) surfaces. Bar = 0.1 mm. **H**. Bottom portion of the flower in Fig. 2e, showing subtending bracts (br, arrows). Bar = 0.5 mm. **I**. An elongated ovule with a micropyle-like depression (arrow) inside the ovary of Flower 9 in Fig. S5e. The inset shows the details of the micropyle-like depression. Bar = 0.5 mm, and inset bar = 0.1 mm.

**Figure 3.**
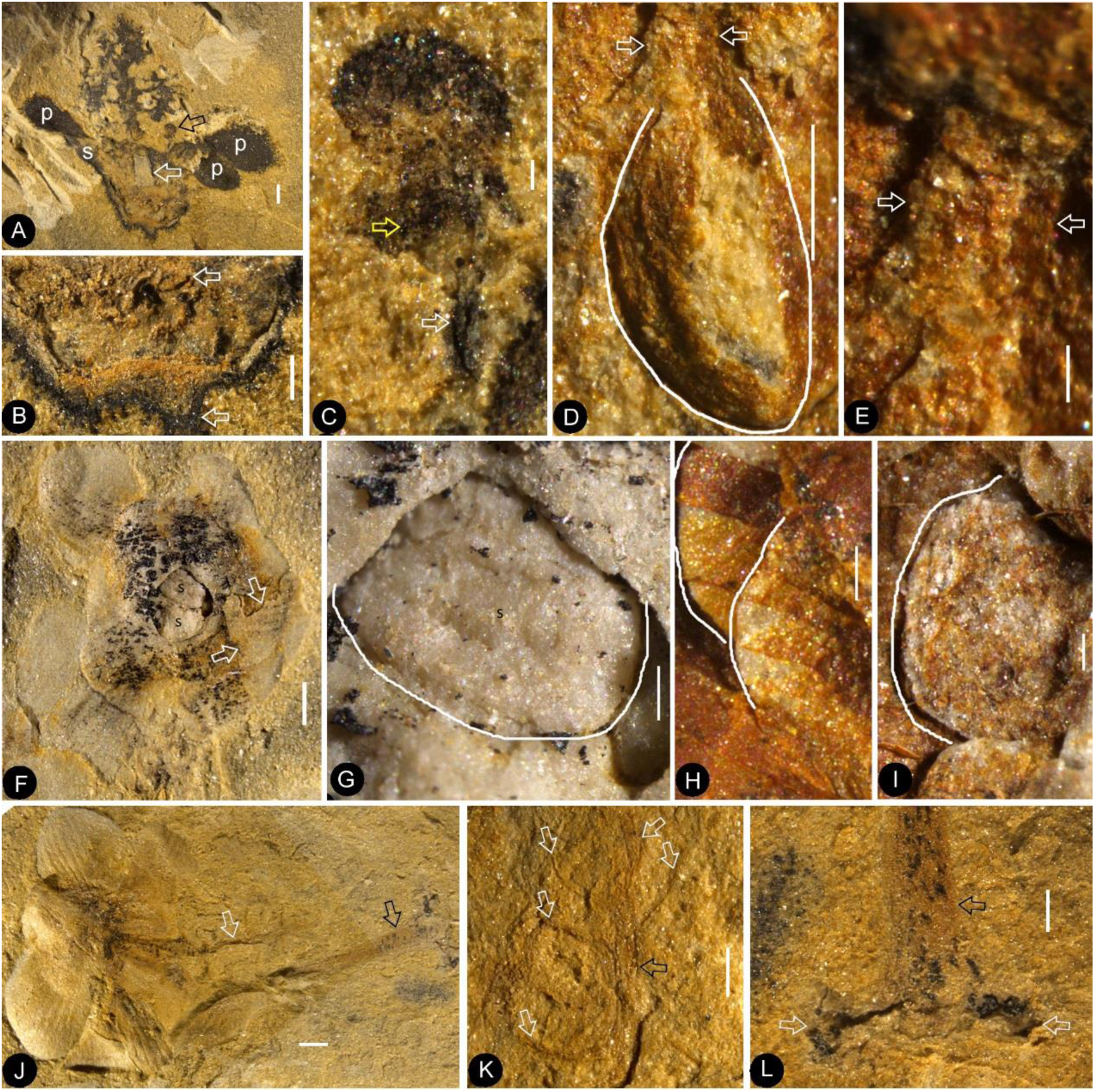
Dendroid style, *in situ* seeds/ovules, and details of flowers. Bar = 1 mm except otherwise annotated. **A**. A longitudinally split flower (counterpart of Flower 10 in Fig. S5e) showing the sepal (s) and petals (p), style base (white arrow), and an unknown organ (black arrow). **B**. Detailed view showing the pedicel (lower arrow) terminating at the bottom of the receptacle in Fig. 3a. Note the ovarian roof (upper arrow). Bar = 0.5 mm. **C**. An unknown organ with a short stalk (white arrow) and enlarged terminal (yellow arrow), enlarged from Fig. 3a. Bar = 0.1 mm. **D**. An ovule (white line) hanging by its funiculus (between arrows) on the ovarian wall of the Flower 2 in Fig. S5b. Bar = 0.5 mm. **E**. Detailed view of the funiculus (between arrows) of the ovule in Fig. 3d. Bar = 0.1 mm. **F**. Top view of Flower 8 in Fig. S5e with sepals and petals surrounding the receptacle/ovary containing two seeds (s). Note the residue of the ovarian roof fragment (arrows). PB22222a. **G**. Detailed view of one of the oval seeds (s) inside the receptacle in Fig. 3f. PB22222a, Bar = 0.2 mm. **H**. Two seeds (white line), one overlapping the other, inside the receptacle shown in Fig. 1f. PB22226. Bar = 0.2 mm. **I**. An oval seed (white line) inside the receptacle/ovary of Flower 7 in Fig. S5e. Bar = 0.2 mm. **J**. Detailed view of flower 17 in Fig. S5e, showing petals (p) with characteristic longitudinal ribs, connected dendroid style (white arrow), and a pedicel (black arrow). **K**. Branches (white arrows) on the main style (black arrow) of the flower in Fig. 3j. Bar = 0.5 mm. **L**. Flower pedicel (black arrow) connected to a branch (white arrows), of the flower in Fig. 3j. Bar = 0.5 mm.

**Figure 4.**
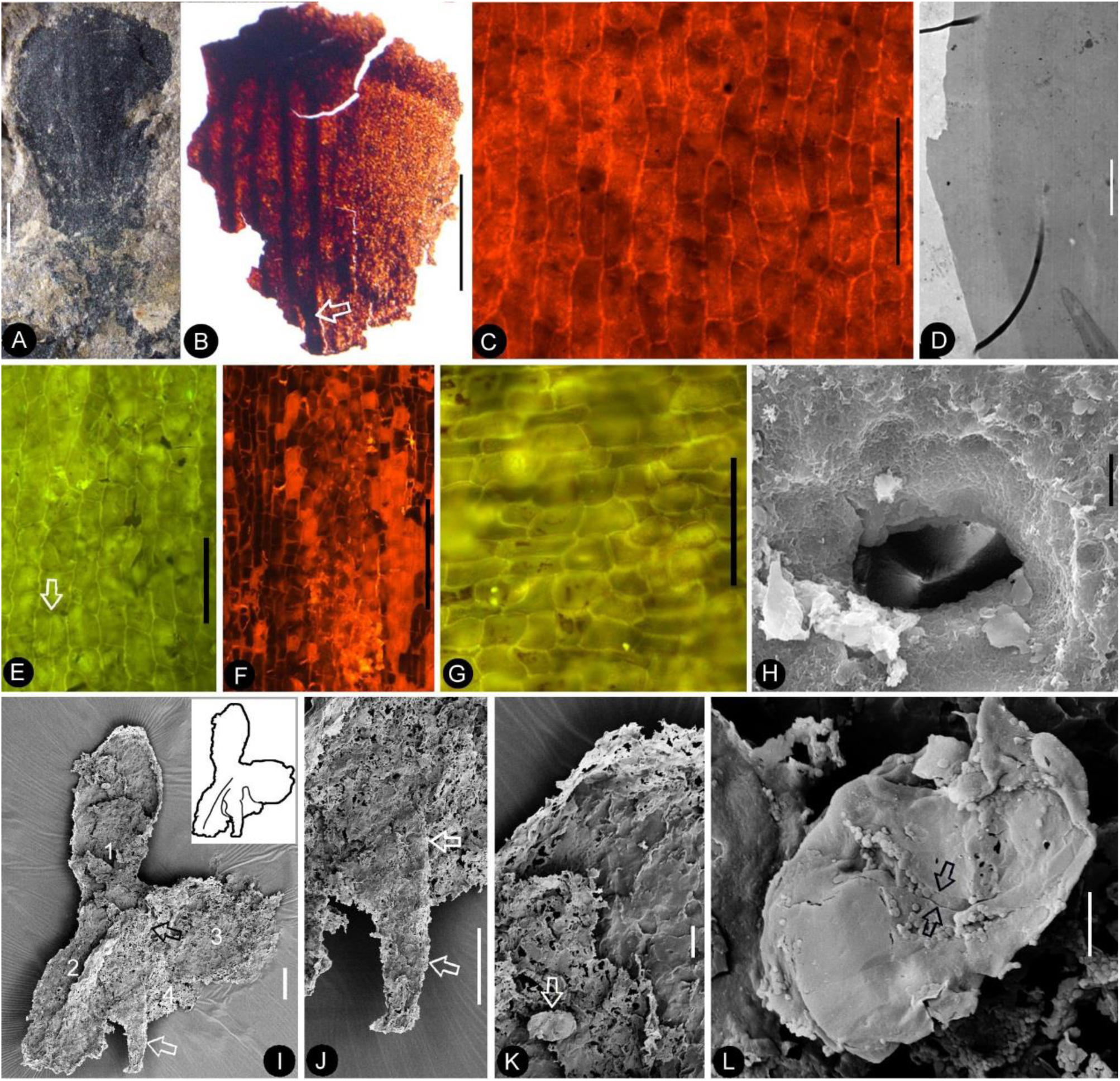
Details of the leaf, sepal, petal, and possible stamen. A-C, stereomicroscopy; C-G, fluorescence light microscopy; H-L, SEM. Bar = 1 mm except otherwise annotated. **A**. A petal with a narrowing base. PB22280. Bar = 1 mm. **B**. A partial petal from Flower 2 in Fig. 1b, with the longitudinal rib (to the left) forking at the base (arrow) and the rib-free wing to the right. **C**. Elongated epidermal cells of the petal in Fig. 4b. Bar = 0.1 mm. **D**. Transmission electron microscope view showing the cuticle (left, light color) of the petal removed from the flower in Figs. S6g-i. PB22223. Bar = 2 μm. **E**. Elongated epidermal cells not in strict longitudinal files in the wing portion of the petal in Fig. 4b. Note the two newly formed epidermal cells (arrow). Bar = 0.1 mm. **F**. Ribs with elongated epidermal cells (left and right) alternating the between region with less elongated cells (middle) of the petal in Fig. 4b. Bar = 0.2 mm. **G**. Elongated (above) and isodiametric (below) epidermal cells on the sepal of Flower 2 in Fig. 1b. Bar = 0.1 mm. **H**. A stoma on the bract of the flower in Fig. 1h. Bar = 5 μm. **I**. Possible stamen under the petal marked by arrows in Fig. S6g-i. Note the short stalk (arrow), conspicuous pollen sacs (1–4), and the position (black arrow) of the possible *in situ* pollen grain shown in Fig. S6j. Bar = 0.1 mm. **J**. Stalk (arrows) of the possible stamen in Fig. 4i. Bar = 0.1 mm. **K**. Detailed view of pollen sac 1 in Fig. 4i. Note a single *in situ* psilate pollen grain (arrow). Bar = 20 μm. **L**. Possibly monocolpate psilate pollen grain with a colpus (between arrows). Bar = 5 μm.

**Figure 5.**
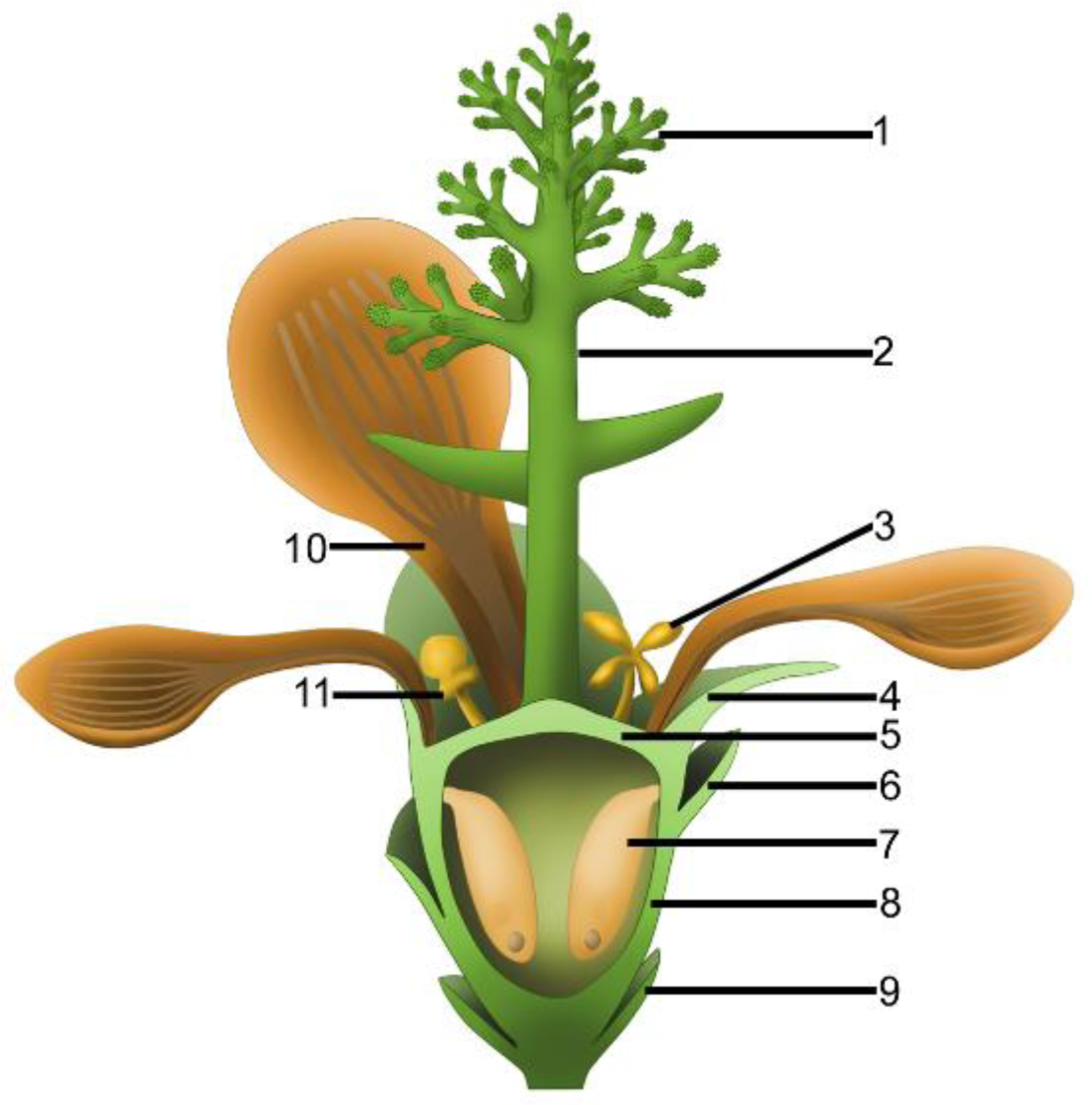
Reconstruction of *Nanjinganthus*. 1, branches of dendroid style; 2, dendroid style; 3, possible stamen; 4, sepal; 5, ovarian roof; 6, scale; 7, ovule; 8, cup-form receptacle; 9, bract; 10, petal; 11, unknown organ.

**Specific diagnosis**: the same as the genus.

**Description**: The flowers are frequently concentrated and preserved in groups on certain bedding surfaces (Figs. 1a-b).

***Flower bud*** A flower bud is preserved as a coalified compression, 6.4 mm long and 3 mm wide, with characteristic longitudinal ribs on the sepals and petals (Fig. 1g). The sepals are 1.3–2.2 mm long and approximately 1.8 mm wide (Fig. 1g). The petals are approximately 3.7 mm long (Fig. 1g). The receptacle/ovary is approximately 3 mm in diameter (Fig. 1g).

***Mature flower*** The flowers are preserved in various states (including coalification), epigynous with an inferior ovary, 8.4–10.7 mm in length and 6.8–12.8 mm in diameter, actinomorphic in the bottom and top views (Figs. 1c-i, 2a, e, 3a,f, S4h, S5a-e, S6a-b, d-i, S7a-b,d,g, S8d-h). The pedicel is approximately 12 mm in length and 0.76 mm in diameter (Figs. 3a, b, j, l). Basally fused bracts are observed at the bottom in a few flowers, and a stoma is seen on a bract (Figs. 2e, h, S7g-h, S9d). Scales are attached on the periphery of the receptacle/ovary (Figs. 1h-i, S6a-b, S7a, g, S8i). The sepals are 1.7–3 mm long and 2.7–4.3 mm wide, with two lateral rib-free wings, with about four longitudinal ribs in the center, and attached to the receptacle rim with their whole bases (Figs. 1f, 2e, S6d-f, S7a, d-e, S8d-e, h-i, l). The epidermal cells are elongated, 44–156 μm x 33–54 μm, with straight cell walls are seen in the middle region, while isodiametric epidermal cells 16–71 μm x 10–54 μm are seen in the lateral wings of the sepals (Figs. 4g, S9e-g, S10h-k). The petals are 3.1–6.6 mm long, 1.9–5.4 mm wide, compressed to only about 11 μm thick, with two lateral rib-free wings, with a cuneate base and longitudinal ribs in the center, inserted inside the sepals and on the rim of the receptacle (Figs. 1c-f, h-j, 2a, e, 3a, 4a, S6a-b, d-f, S7a-b, d, f-g, S8f-i, k, S10a-b, d-e). The ribs are approximately 0.12 mm wide, forking only basally, with elongated epidermal cells with straight cell walls, 32–144 μm x 17–30 μm on the abaxial and 19–72 μm x 13–29 μm on the adaxial (Figs. 4a-f, S9a-b). The lateral wings are free of ribs, and each is approximately 1.2 mm wide, with more or less isodiametric epidermal cells 23–64 μm x 18–37μm (Figs. 4a, e, S10g). A possibly immature stoma is seen on one of the petals (Fig. S9c). A possible stamen is at least 1.1 mm long and 0.8 mm wide, with a stalk (equivalent to filament) at least 0.23 mm long and 43–76 μm in diameter (Figs. 4i-j). Four pollen sacs terminate the stalk, each 0.53 mm long and 0.27 mm wide (Figs. 4i-j). Possible *in situ* pollen grain is about 30 × 20 μm, psilate and possibly monocolpate (Figs. 4k-l, S6j). The receptacle is cup-form, 3–4.8 mm in diameter and 2–4.5 mm high, surrounded by a 0.3 mm thick wall in the bottom and sides, and covered by an ovarian roof about 0.2 mm thick from the top (Figs. 1h-i, 2a, e-h, 3a-b, f, S8d, f-g). The ovarian roof is horizontal, with smooth integral outer and inner surfaces, 0.14–0.22 mm thick, with a style vertically inserted on its center (Figs. 2a-c, e-g, S7c). The style is up to 1 mm in diameter, with lateral branches adding 2.6–4.4 mm width to the style (Figs. 1h-i, 2a-d, 3a, j-k, S6a-c, S8i-j). Each ovary contains two or more ovules that are 0.65–3 mm × 0.5–1.7 mm, elongated or oval-shaped, hanging on a 0.08–0.27 mm wide funiculus inserted on the inner wall of the receptacle/ovary (Figs. 1f, 2i, 3d-i, S5e, S8a-e). A micropyle-like depression 0.1–0.19 × 0.19–0.25 mm is seen on an ovule (Figs. 2i, S5e, S8a).

**Holotype**: PB22222 (Figs. 1a, 1c-d, 3f-g, S5e, S6c).

**Isotypes**: PB22221, PB22223-PB22229, PB22236, PB22238, PB22241-PB22243, PB22245-PB22247, PB22256-PB22260, PB22278-PB22282, PB22489.

**Etymology**: *dendrostyla*, for “tree-like” (*dendri*-) and “style” (-*stylus*) in Latin.

## Discussions

Several gymnosperm groups, including Caytoniales, Corystospermales, Ginkgoales, Czekanowskiales, Iraniales, Pentoxyales, Bennettitales, Coniferales, and Gnetales, flourished during the Mesozoic. However, these taxa have been well-studied and understood. Knowledge of these plants indicates clearly that *Nanjinganthus* distinguishes itself from all known gymnosperms, fossil and extant, by its probable bisexuality and with enclosed ovules in invaginated receptacle (see **Supporting Materials and Table 1**). The enclosed ovules makes it out of the question whether *Nanjinganthus* stands for another extinct gymnosperm, just as what happened to Caytoniales (Thomas 1925; Harris 1940; Barbacka and Boka 2000).

**Table 1.** Comparison between *Nanjinganthus* and Mesozoic gymnosperms.

|  | *Nanjinganthus* | *Caytonia* | Bennettitales | Corytospermales | Ginkgoales | Coniferales | Iraniales | Czekanowskiales | Pentoxiales |
| --- | --- | --- | --- | --- | --- | --- | --- | --- | --- |
| Symmetry | Radial | Bilateral | Radial | Bilateral | Radial | Radial | Radial? | Bilateral | Radial? |
| With foliar appendages | Yes | No | No | No | No | ? | No | No | No |
| Enclosed ovule | Yes | No | No | No | No | No | ? | No | No |
| Opening in female part | No | Adaxial basal | ? | Adaxial basal | N/A | N/A | ? | Distal slit | ? |
| Penetrating cone axis | No | ? | Yes | ? | ? | Yes | ? | Yes? | Yes |

Although multiple characters have been suggested to identify fossil angiosperms (Herendeen *et al*. 2017), angio-ovuly before pollination is the only character that guarantees an angiospermous affinity (Tomlinson and Takaso 2002; Wang 2010). This criterion has been repeatedly applied to identify fossil angiosperms (i.e. *Archaefructus* (Sun *et al*. 1998)), because, at least initially, no other features (stamen, venation, pollen grains) were available to support its angiospermous affinity. In our material an integral ovarian roof has no any opening (Figs. 1h-i, 2e-g, 3a-b,f, S7c, and S8c,f-h). In some specimens this ovarian roof blocked the sediment from entering the ovarian locule (Figs. 2f-g). This suggests a full enclosure of ovules, confirming the angiospermous affinity for *Nanjinganthus*.

The radial arrangement of two whorls of foliar parts (sepals and petals) in *Nanjinganthus* is very similar to those of flowers in extant angiosperms (Figs. S11a-b). A closed ovary, 4-parted anther, probably bisexual reproductive organ, and low number of ovules/seeds seen in *Nanjinganthus* (a character combination never seen in any known gymnosperm) emphasizes that *Nanjinganthus* cannot be interpreted as a gymnosperm. Furthermore, *Nanjinganthus* satisfies all thirteen definitions of flowers advanced by various authors (Bateman *et al*. 2006). These features reinforce the angiospermous affinity of *Nanjinganthus* (Fig. 5).

There have been several suggested models of ancestral angiosperms (Arber and Parkin 1907; Cronquist 1988; Endress and Doyle 2015; Sauquet *et al*. 2017). These models were drawn more or less after the assumed basalmost living angiosperms, *Magnolia* or *Amborella*. The common features of these model plants include apocarpy, superior ovary, lack of obvious style, etc. However, none of these features are seen in *Nanjinganthus*. Instead, an inferior ovary is clearly seen in *Nanjinganthus*, a feature unexpected by most theories of angiosperm evolution. This discrepancy suggests EITHER that inferences based on living plants have limited capability of “predicting” past history, OR that angiosperms originated polyphyletically and each lineage has followed a different evolution route, OR that angiosperms have a history longer than the Cretaceous, OR a combination of these. Whatever the implications are, the dominant theories of angiosperm evolution apparently do not reflect the botanical reality.

Most *Nanjinganthus* specimens are concentrated to a limited number of bedding surfaces, and up to 82 individuals are preserved on a single slab (Figs. 1a-b, S5a-g), suggesting that *Nanjinganthus* may have flourished and dominated a certain niche. The concentrated preservation of delicate flowers is more likely a result of autochthonous preservation, suggesting a habitat very close to water for *Nanjinganthus*.

Various studies including palaeobotany of the South Xiangshan Formation in the last century (Hsieh 1928; Li *et al*. 1935; Sze and Chow 1962; Zhou and Li 1980; Cao 1982; Wang *et al*. 1982; Huang 1983; Ju 1987; Huang 1988; Cao 1998; 2000) and our palynological analysis and U/Pb dating (Figs. S1b, S1f-g; Table S2; (Santos *et al*. in progress)) suggest a latest Early Jurassic age for *Nanjinganthus*. Together with the “unexpectedly” great diversity of angiosperms in the Early Cretaceous (Sun *et al*. 1998; Sun *et al*. 2001; Sun *et al*. 2002; Leng and Friis 2003; Ji *et al*. 2004; Leng and Friis 2006; Wang and Zheng 2009; Wang and Han 2011; Wang and Zheng 2012; Han *et al*. 2013; Wang 2015; Han *et al*. 2017), pollen grains indistinguishable from angiosperms in the Triassic (Hochuli and Feist-Burkhardt 2004; Hochuli and Feist-Burkhardt 2013), a bisexual flower from the Jurassic (Liu and Wang 2016), and an herbaceous angiosperm from the Middle Jurassic (Han *et al*. 2016), *Nanjinganthus* with over 200 specimens is an overwhelming evidence for a pre-Cretaceous origin of angiosperms, fully refuting the recently published self-conflicting review on early angiosperms (Herendeen *et al*. 2017) that does not reflect the true history of angiosperms (Wang 2017).

The systematic position of *Nanjinganthus* is apparently open to further investigation, although it demonstrates certain resemblance to Pentapetalae *sensu* (Judd et al. 2016). We cannot determine whether *Nanjinganthus* stands for a Jurassic stem group of angiosperms that started their radiation in the Cretaceous or a lateral branch leading to dead end of evolution. It is premature to determine its phylogenetic position before more information of contemporaneous plants is available, although we welcome other phylogeneticists to evaluate *Nanjinganthus* in their own ways and perspective.

## Conclusion

The recognition of *Nanjinganthus* is based on at least 284 individual flowers from the Early Jurassic that are preserved in various states and orientations. The enclosed ovules suggest an angiospermous affinity for *Nanjinganthus*. The discovery of *Nanjinganthus* will trigger a re-thinking on the origin and early history of angiosperms and call for modifications to the current theories of angiosperm evolution.

## Acknowledgements

We thank Ms. Chunzhao Wang for help with SEM during this research, and Dr. Walter Judd at the University of Florida for comments and suggestions. This research was supported by National Natural Science Foundation of China (41688103, 91514302, 41572046), Strategic Priority Research Program (B) of Chinese Academy of Sciences (Grant No. XDPB05) awarded to X.W.; and State Forestry Administration of China (No. 2005–122), Science and Technology Project of Guangdong (No. 2011B060400011), and Special Funds for Environmental Projects of Shenzhen (No. 2013–02) awarded to Z. J. L.. This is a contribution to UNESCO IGCP632. We declare no competing interests.

## Author Contributions

QF, XW collected the fossil materials and initaited the study; MP did the cuticle preparation; XW performed the sample processing, photography, microscopy, SEM and TEM and observation; QF, JBD, MP, ZJL, QZ, XW did botanical analysis and drafted the manuscript; MGÁ did the reconstruction drawing; YH, PY did the Micro-CL; HC did the isotopic dating; JBD did the palynological dating; KD did the TEM sample preparation and observation. ALL finalized the manuscript.

